# DGINN, an automated and highly-flexible pipeline for the Detection of Genetic INNovations on protein-coding genes

**DOI:** 10.1101/2020.02.25.964155

**Authors:** Lea Picard, Quentin Ganivet, Omran Allatif, Andrea Cimarelli, Laurent Guéguen, Lucie Etienne

## Abstract

Adaptive evolution has shaped major biological processes. Finding the protein-coding genes and the sites that have been subjected to adaptation during evolutionary time is a major endeavor. However, very few methods fully automate the identification of positively selected genes, and widespread sources of genetic innovations as gene duplication and recombination are absent from most pipelines. Here, we developed DGINN, a highly-flexible and public pipeline to Detect Genetic INNovations and adaptive evolution in protein-coding genes. DGINN automates, from a gene’s sequence, all steps of the evolutionary analyses necessary to detect the aforementioned innovations, including the search for homologues in databases, assignation of orthology groups, identification of duplication and recombination events, as well as detection of positive selection using five different methods to increase precision and ranking of genes when a large panel is analyzed. DGINN was validated on nineteen genes with previously-characterized evolutionary histories in primates, including some engaged in host-pathogen arms-races. The results obtained with DGINN confirm and also expand results from the literature, establishing DGINN as an efficient tool to automatically detect genetic innovations and adaptive evolution in diverse datasets, from the user’s gene of interest to a large gene list in any species range.

## Introduction

Genetic innovation is a major adaptation process that has impacted genome structures and functions over millions of years in response to natural selection. Such changes have shaped key biological functions, such as reproduction, adaptation to a new environment, immunity, sensory-perception, host-pathogen interaction. Adaptation in protein-coding genes can take place through several mechanisms. They include, amongst others, positive selection on coding sequences, duplication events with subsequent divergence of the copies, as well as recombination (Daugherty and Malik 2012). The first is caused by natural selection that increases the frequency of advantageous mutations, leading to an apparent excess of non-synonymous substitution rates over synonymous ones over evolutionary times. This notably leads to the accumulation of beneficial amino-acid changes at the location of functionally important residues, such as the interface of proteins involved in host-virus interactions. Gene duplication is another important source of genetic novelty, which notably allows to increase the general evolvability (Daugherty and Zanders 2019, Kondrashov 2012). The fixation of multiple copies enables diversification of gene function through subfunctionalization or neofunctionalization. Moreover, gene conversion, by recombination between alleles, allows for rapid divergence of the copies. Gene duplication and loss may further be a dynamic and rapid adaptation process (McLaughlin and Malik, 2017, Daugherty and Zanders 2019, Kondrashov 2012).

These mechanisms fueling genetic novelty are all parts of the response of organisms to selective pressures and must therefore be analyzed as much has possible together to wholly apprehend the evolutionary history of genes. However, despite their frequency and their biological importance and relevance, these diverse evolutionary innovations are not accounted for in most tools and studies analyzing genes under adaptive evolution (such as Kosiol et al., 2008, Hawkins et al 2019, and reviewed in Sahm et al 2017). Lastly, performing gold-standard and complete phylogenetic analyses is usually highly hand-curated. Our goal was therefore to design a tool that would incorporate all these mechanisms at the origin of genetic innovation in a robust end-to-end pipeline to identify and characterize new protein-coding genes with signatures of adaptive evolution.

Such a pipeline requires the automation of essential steps. Primarily, searching for homologous gene sequences and identifying orthologous relationships represent a time-consuming and difficult process. No existing tool include these steps, because they either remain essentially hand-curated (Hyphy suite (Pond et al., 2005), Selecton (Stern et al., 2007), IDEA (Egan et al., 2008), JcoDa (Steinway et al., 2010), PoSeiDon (Fuchs et al., 2017) and POTION (Hongo et al., 2015)), are restricted to specific vertebrate and prokaryotic species (PhyleasProg (Busset et al., 2011) and PSP (Su et al., 2013)), or rely on published orthologous annotations (essentially from the NCBI HomoloGene) which may become imprecise on non-model species.

Secondly, correct codon alignments are necessary for the accurate detection of residues under positive selection. However, current pipelines rely on protein or nucleotide alignment softwares like ClustalW (Thompson et al., 1994) or Muscle (Edgar, 2004), although more recent ones such as PRANK (Löytynoja and Goldman, 2008) have been repeatedly shown to provide high-quality codon alignments, thereby diminishing false positives during the detection of positive selection (Fletcher and Wang, 2010, Privman et al., 2012, Jordan and Goldman, 2012, Markova-Raina and Petrov, 2011).

Thirdly, we identified the need to include within a single analysis the detection of positive selection signatures by different methods and models, to allow for more specificity and sensitivity of the results, as well as to help “ranking” genes in an evolutionary screening approach (for example Abdul et al., 2018, Elde et al 2009, Schultz and Sackton 2019, Malfavon-Borja et al., 2013, McBee et al., 2015, Rowley et al., 2016). Moreover, the inclusion of methods in which the experienced user has access to the parameterization of the maximum likelihood models is needed (van der Lee et al, 2017). Existing tools rely almost exclusively on PAML codeml (Yang, 2007), which has allowed the identification of numerous genes under positive selection, but offers limited options for parameterization.

Overall, there seemed to exist a void when it comes to pipelines which fully automate the search for adaptive evolution in protein-coding genes, from retrieving homologous sequences of a gene of interest in any species range, establishing orthologous relationships, reconstructing codon alignments and the corresponding phylogenies, to detecting different genetic innovations using gold-standard and diverse methods to ensure high-degree of confidence in the results. We thus developed an integrative pipeline, that we named DGINN (for Detection of Genetic INNovations) to satisfy those requirements. All scripts are freely available on Github and as a docker on DockerHub. We also focused on user-friendliness and flexibility, so that biologists can use with ease and use only parts of the workflow for various purposes. DGINN was developed as a one-gene workflow and can easily be up-scaled to screen large datasets of dozens or hundreds of genes. Finally, we performed an extensive validation of our pipeline, using published and highly hand-curated phylogenetic data on a set of nineteen primate genes with various evolutionary histories including genes involved in virus-host evolutionary arms-races (Daugherty and Malik 2012, Duggal and Emerman 2012). Through DGINN, we further identified previously uncharacterized signatures of genetic conflict in the primate Guanylate-binding protein (GBP) family, which plays important roles in cell-autonomous immunity against pathogens (Kim et al 2012, Kraap et al., 2016).

## Materials and Methods

### Pipeline structure

The overall goal of the DGINN pipeline (overviewed in Figure 1) is to provide an easy, integrated, and robust way of detecting genetic innovations from a gene sequence provided by the user on two scales, either on one specific gene for fine-tuned analyses or on large sets of genes of interest for screening purposes.

**Figure 1.**
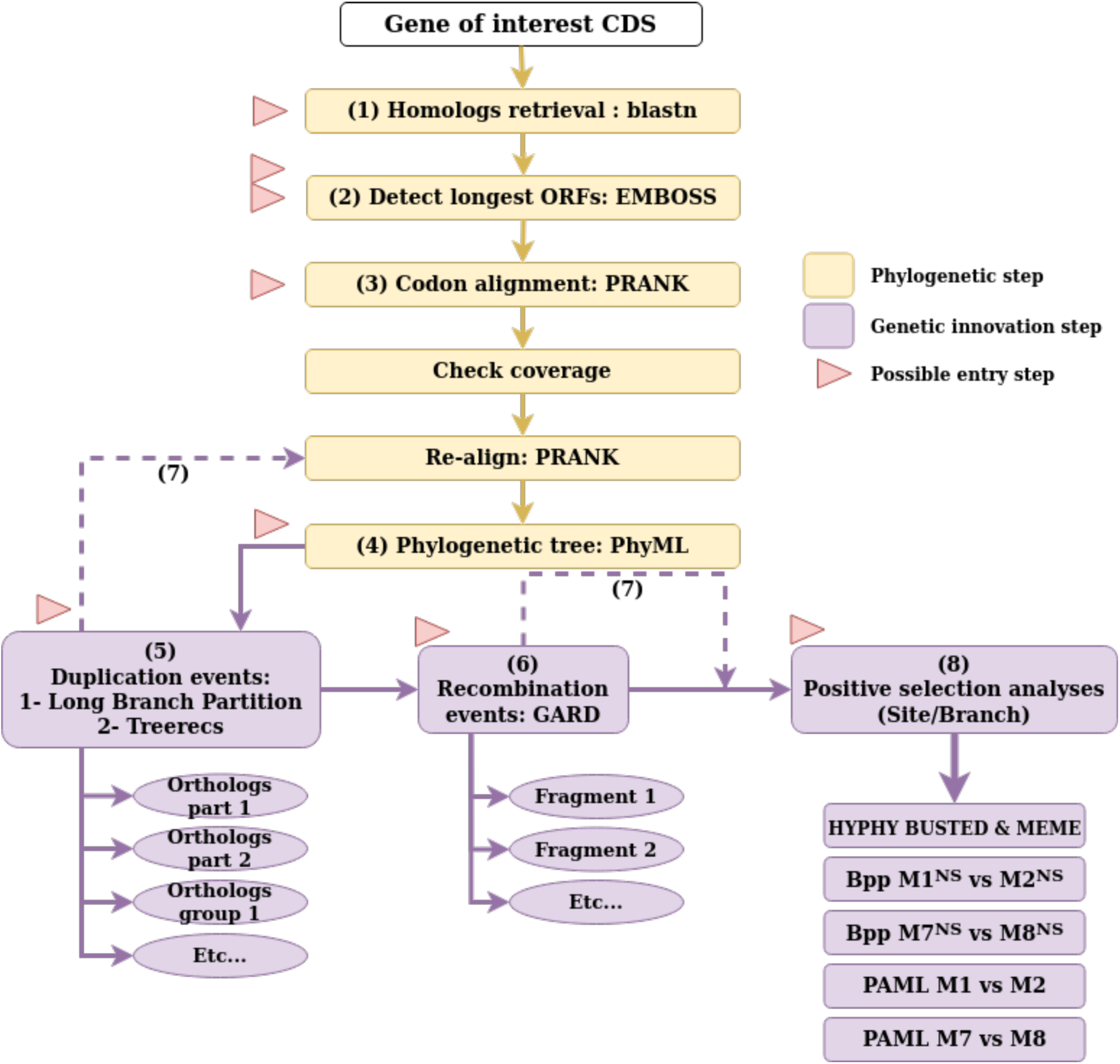
Workflow diagram of DGINN. Phylogenetic steps (yellow) happen sequentially from the entry point of the pipeline (Steps 1-4). Each genetic innovation step (purple, Step 5, 6, and 7) is optional. All red arrowheads denote possible entry points into the pipeline following file formats from Table 1.

DGINN is implemented in Python and uses numerous modules, including some from Biopython, as well as several independent softwares. The list of modules and external softwares is provided in the pipeline documentation. All scripts and documentation can be downloaded from Github. To enhance user-friendliness, options are handled through a parameter file, minimizing the complexity of the command line. Importantly, a Docker image is also available for local use without manual installation of the external required softwares. The Docker may also be used to screen large dataset using AWS Batch for example (https://aws.amazon.com/batch/). A specific script for the extraction of batch results, parseResults.py, and a graphical interface to produce basic figures with them, have also been developed (see Availability).

**Table 1.** Overview of the possible entry steps into DGINN. DGINN can be entered at different steps to enhance flexibility. If the user introduces the name of the proper entry step option and inputs the appropriate files for this option in the parameter file, DGINN will start at that step, ignoring the upstream steps.

| Name of entry step option | Input files | Format |
| --- | --- | --- |
| blast | CDS of the gene of interest | Fasta |
| accession | List of blast results | NCBI tabulated format |
| fasta | List of accession identifiers (one per line) | Text file |
| orf | mRNA sequences of homologs | Fasta |
| alignment | CDS sequences of homologs/orthologs | Fasta |
| tree | (codon) alignment of homologs/orthologs | Fasta |
| duplication | (codon) alignment, gene tree | Fasta, newick |
| recombination | (codon) alignment | Fasta |
| positiveSelection | codon alignment, gene tree | Fasta, newick |

The overall workflow of the DGINN pipeline is a succession of eight steps, described hereafter. Of note, DGINN is designed to be extremely flexible as to its uses. The user can enter the workflow at any step with the files resulting from their own analyses, as indicated in Table 1 and Figure 1. The name of the step reflects the very first step performed with the option. For example, starting DGINN at the ‘blast’ step will make it begin with the blast search, and then execute the whole pipeline. The duplication, recombination and positive selection steps will not be performed if the user has not specifically opted in for them, allowing for maximum flexibility.

#### (Step 1) Automated retrieval of homologous genes in species of interest

DGINN uses BLAST+ search (Camacho et al., 2009) against the NCBI databases. The BLAST search can be done against a local database constructed by the user, or online against specific NCBI databases, which allows the user to limit the search to certain sequences, such as ESTs, or certain species, by providing the proper Entry Query, following the syntax used on the NCBI website, as described in their documentation (https://www.ncbi.nlm.nih.gov/books/NBK3837/#EntrezHelp.Entrez_Searching_Options). BLAST+ is used by providing the coding sequence of the gene of interest against a nucleotide databank (blastn). We decided not to use blastp (protein query against protein database) as it significantly complicated the recuperation of the nucleotide sequences afterwards, which are indispensable to the rest of the pipeline. Moreover, nucleotide databases include more sequences and thus allow for a more exhaustive search. The number and speed of requests against NCBI databases can be increased through the acquisition of an NCBI API key, available online. This ensures access to the largest possible number of sequences, including those not annotated as orthologous or paralogous sequences. The user may modify minimum e-value, coverage, and identity values to reflect the specificities of the database and the species set against which they are using BLAST+. Because we validated our pipeline on primate evolution, we set those with default values of 10^−4^, 50%, and 70%, respectively, to retrieve a maximum of homologous sequences without too many unrelated sequences.

#### (Step 2) Elimination of overly long sequences and isolation of Open Reading Frames (ORFs)

Because the user may want to cast a wide net in terms of homologue retrieval, and thus use low coverage and identity for the blastn search (Step 1), a variety of resulting hits are retrieved, including overly long sequences from whole contigs or chromosomes. Those sequences considerably increase the analysis time if not properly curated. Furthermore, the detection of ORFs of interest is extremely difficult, as they contain numerous genes. In DGINN, we identify and remove such sequences based on the median length of all the retrieved sequences: if the median is longer than 10,000 nucleotides, any sequence longer than twice the median are taken out, otherwise sequences are deleted if they exceed three times the median length. The remaining sequences are searched for ORFs using ORFinder from the EMBOSS package (Rice et al., 2000) to keep only the coding sequence of each gene. The longest detected ORF of each sequence is selected for further analysis.

#### (Step 3) Initial codon alignment

Positive selection analyses rely on identifying substitutions leading to amino-acid changes over those being silent. Therefore, a codon alignment of good quality is essential. However, very few softwares propose true codon-alignment modes. To date, the best codon aligners are PRANK (Löytynoja and Goldman, 2008) and MACSE (Ranwez et al., 2011). PRANK has been shown to produce the best alignments for positive selection analyses (Schneider et al., 2009, Fletcher and Yang, 2010, Markova-Raina and Petrov, 2011, Jordan and Goldman, 2012, Privman et al., 2012). From our observations, MACSE also produced high-quality codon alignments, but it was significantly slower than PRANK. We therefore selected the latter as the best solution for both quality alignments and lower computational time. PRANK alignments are performed with the codon model and without forcing insertions to be skipped, and otherwise default settings (prank -F - codon; version 150803). After this initial alignment, we added a quality control step to eliminate sequences that did not align properly, using Python homemade scripts, based on alignment coverage against the query (either the user-provided value or default of 50%). The remaining sequences are then re-aligned using the same settings.

#### (Step 4) Construction of the initial phylogenetic gene tree

The gene’s phylogenetic reconstruction is performed with PhyML v3.2 (Guindon et al., 2010). We opted for a HKY+G+I model as default, because it offers the best combination of realistic phylogenies without being too time-consuming. As the produced trees are only intended for screening purposes at this step, we also opted to use approximate Likelihood Ratio Test (aLRT) for the statistical support of the branches (Anisimova and Gascuel, 2006).

#### (Step 5) Identification of duplication events and orthologous groups

As previous steps retrieved homologues without relying on synteny or gene annotation, we implemented two strategies to identify duplicated genes and to constitute orthologous groups necessary for the positive selection analyses. DGINN first identifies the overly “long branches” within the gene tree. We define a “long branch” as a branch which length is superior to 50 times the mean of all branch lengths in the tree (i.e. the estimated number of substitutions per position is at least 50 times superior in the “long branch” compared to the mean). When “long branches” are identified, the tree is cut along those “long branches” and the groups of sequences subsequently constituted are re-aligned (back to step 3) and their trees recomputed separately (step 4). This constitutes a first method of separating highly divergent groups of genes, between which detection of positive selection may be ambiguous because of suspicion of paralogy and branch length saturation. However, for multigenic families that include paralogues that have recently diverged, the gene members cannot be separated solely based on the relative lengths of the tree branches. We therefore included a phylogenetic reconciliation method, TreeRecs (Comte et al., 2019), to identify genes sharing a common evolutionary history in our species of interest. To identify duplication events, TreeRecs reconciles a user-provided species tree or cladogram to each gene tree. From the reconciled tree, DGINN establishes groups of orthologues based on ancestral duplication events annotated on the reconciled tree. Since interspecific positive selection analyses rely on the comparison of several orthologous sequences, orthologous groups resulting from very recent duplications may have too few sequences to be informative for those analyses. So we chose to ignore duplication events that were not ancestral enough, by taking into account the minimal number of species represented downstream of the event. This number is user-determined. We decided on a default setting of a minimum of eight species to extract a duplication group from the original alignment, based on the results obtained by Anisimova et al. (2002), and in primates specifically by McBee et al., (2015). Duplication events on nodes that do not have at least two species in common in the groups formed on either side of the node are considered dubious: the corresponding annotated events are then ignored by DGINN. After extraction based on ancestral duplication events, the orthologous groups are realigned using PRANK as in Step 3.

#### (Step 6) Identification of recombination events and splitting of alignments along the significant breakpoints

To account for recombination, DGINN includes GARD from HYPHY (Kosakovsky Pond et al., 2006) with standard parameters. The breakpoints are then assessed for statistical significance using a likelihood ratio test (LRT) with p < 0.05 against a null hypothesis that there is no breakpoint at that position. If any breakpoint is significant, it is moved to the nearest inter-codon site, and the alignment is subsequently cut into the corresponding non-recombinant fragments. These non-recombinant alignments, as well as the original one, will become the input in the following steps.

#### (Step 7) Construction of the final phylogenetic trees

Following the analyses of duplication and recombination events (steps 5-6), new codon-wise alignments using PRANK (same parameters as in step 3) and new phylogenies using PhyML (same parameters as in step 4) are built for groups of non-recombinant fragments (see step 6) of orthologous genes (see step 5). These final codon alignments and gene trees will further provide the input for the positive selection analyses.

#### (Step 8) Positive selection analyses

Numerous softwares exist to identify positive selection on coding sequences. DGINN includes several methods of positive selection analyses, which the user can chose to turn on or off independently. Those analyses make extensive use of three packages: HYPHY (Pond et al., 2005), PAML codeml (Yang, 2007) through the ETE toolkit (http://etetoolkit.org/), and Bio++ (Guéguen et al., 2013).

From the HYPHY package, we included two methods. First, we included BUSTED (Branch-Site Unrestricted Statistical Test for Episodic Diversification), a random effect model which allows for gene-wide detection of episodic positive selection (Murrel et al., 2015). Results are considered positive in the DGINN pipeline for a p-value < 0.05 for the LRT of the models admitting *vs* not admitting positive selection. Second, we included MEME (Mixed Effects Model of Evolution), which detects individual sites subjected to episodic positive selection based on a mixed effects model (Murrel et al., 2012). These models are complementary, as BUSTED evaluates positive selection at the gene level and MEME at the site level.

Contrary to BUSTED and MEME, the codon substitution models used in PAML codeml focus on pervasive positive selection and not episodic events. Briefly, the codon alignments are fitted to models that do not allow for positive selection, M1 (with two classes ω < 1 and ω = 1) or M7 (where the ω < 1 class is modeled as a gamma law of n classes, n=5 as default in DGINN), and the corresponding models allowing for positive selection with one class of ω > 1 (M2 or M8, respectively). Statistical significance of positive selection is determined through a chi-squared test of the LRT of both associated models (M1 *vs* M2, and M7 *vs* M8) to derive p-values. Results are considered positive in the DGINN pipeline for a p-value < 0.05.

However, PAML codeml relies on the assumption of stationarity (i.e. that the base composition of sequences is at the equilibrium of the evolutionary process), which may impact the detection of selection (Guéguen and Duret, 2018). It is also limited with regards to its parameterization. Therefore, we also integrated the parameterizable Bio++ library to propose similar models but without stationarity assumption (Bio++ models M1^NS^ *vs* M2^NS^, and M7^NS^ *vs* M8^NS^). Similarly, DGINN considers significant positive selection if p-value < 0.05 of each model comparison.

If positive selection is determined with PAML or Bio++, the pipeline will proceed to the identification of the sites under positive selection, using the Bayes Empirical Bayes statistics (BEB) from the M2 and M8 in PAML codeml and the Bayesian Posterior Probabilities (PP) from the M2^NS^ and M8^NS^ models in Bio++. Sites are considered as under significant positive selection if BEB or PP > 0.95.

To detect specific branches/lineages under positive selection, DGINN uses Bio++ to include a method similar to the Free-Ratio test available in PAML codeml, called One Per Branch in DGINN (OPB). The ω ratio is calculated along the branches of the phylogenetic tree by using a M0 model where all parameters but ω are homogeneous. As this step is independent and the Bio++ parameter file is fully accessible, an experienced user can choose any model they wish, allowing for maximum flexibility.

### Pipeline parallelization

DGINN has been developed with the intention to analyze each gene independently, with parallelization over large datasets being handled in a cluster environment. This is done through user-made scripts (such as job arrays) and facilitated through configuration parameters that are specific to this use. -i/--infile allows for easier parallelization by eliminating the need to create parameter files for each analyzed gene. -host/-- hostfile allows the user to indicate the cluster hostfile to avoid conflicts when starting mpi processes.

Also, if the query genes are from human, a separate script is provided for downloading their CCDS sequences prior to using DGINN itself. This script, called CCDSquery.py and available on the Github, only requires a table as its entry, with HUGO Gene Nomenclature Committee (HGNC) approved symbols in one column and the corresponding CCDS accessions in another. This table can be obtained through the HGNC biomart (http://biomart.genenames.org/).

### Results extraction

An independent script, parseResults.py, is provided to extract the essential results after running the pipeline. This script outputs a table (described in DGINN’s documentation) which compiles, for each analyzed gene, the results regarding duplication and recombination events, and the different methods of positive selection detection used (including significance of each method and sites identified). This script only requires the path to the directory containing DGINN’s results as input.

An R Shiny App (see Availability) has been further designed to help the user visualize the results quickly, which only necessitates the file produced by parseResults.py. This app will output the figures in the same format as those shown in Figures 3-4.

**Figure 2.**
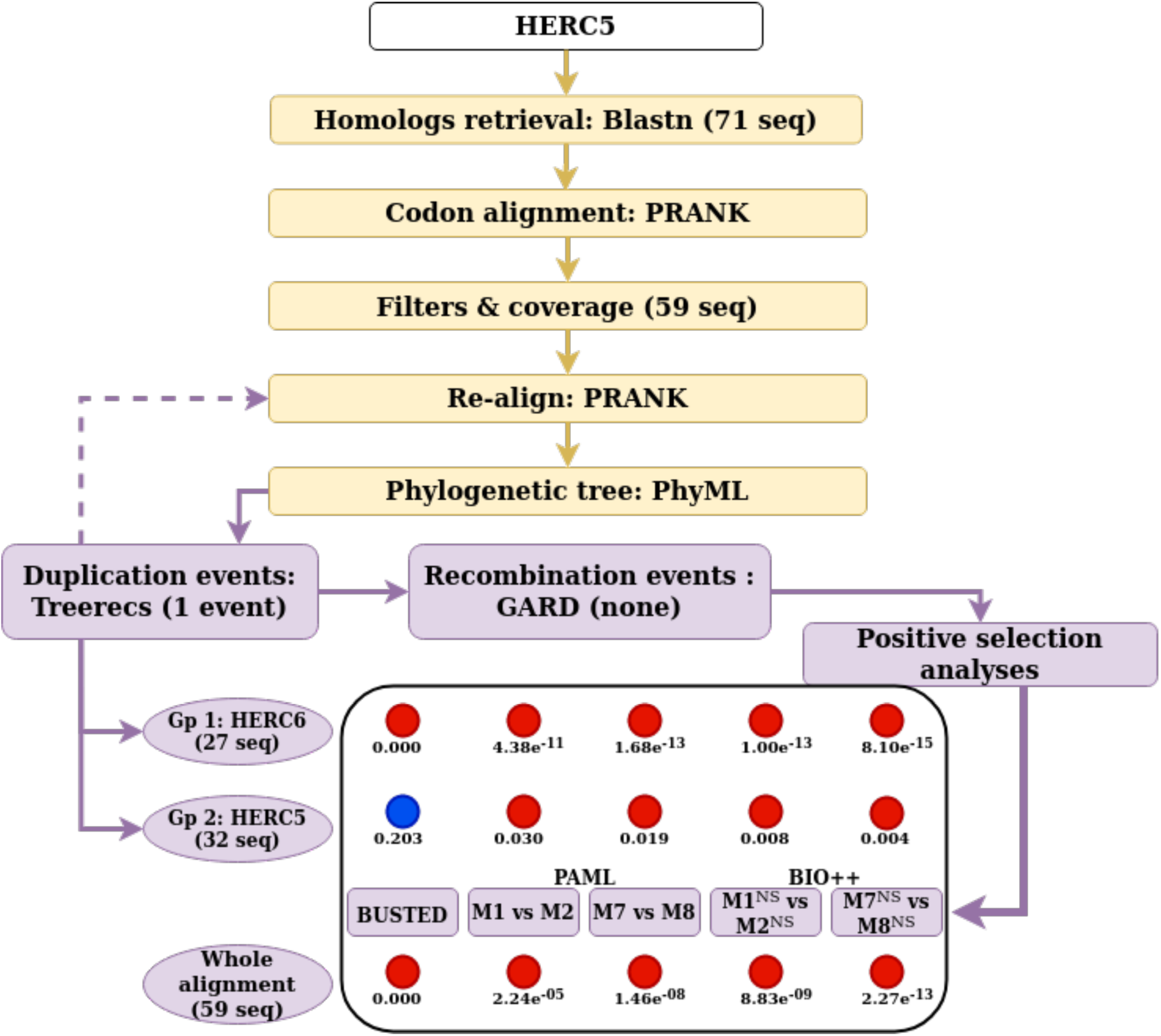
Example of workflow on the HERC5 primate gene. The workflow follows the diagram from Figure 1. Using human HERC5 CDS as the starting point in DGINN gave results for both HERC5 and HERC6. The number of sequences (seq) retrieved or left after each step is indicated. In the bottom panel, each colored circle represents the results from one of the five methods to detect positive selection at the gene level, with red representing significant evidence of positive selection and blue no significant evidence. P-values are indicated below the colored circles. Gp, orthologous group.

**Figure 3.**
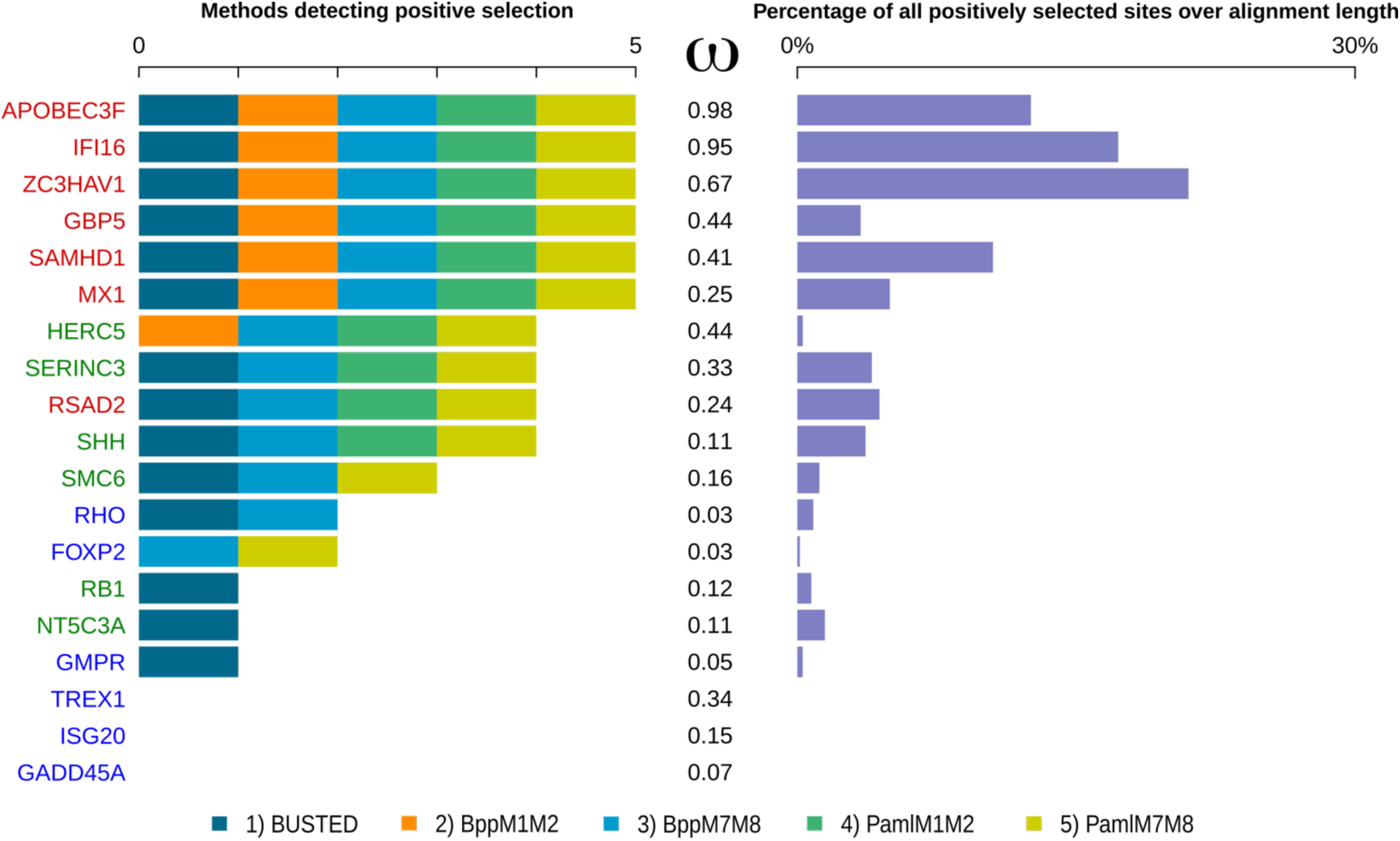
DGINN results on the validation dataset. The nineteen primate genes studied are color-coded according to their selection profile category (Table 2). Left panel, number of methods detecting significant positive selection for each alignment; each method is color-coded (embedded legend). Middle, mean ω value calculated by Bio++ M0 model. Genes are ordered by descending number of methods detecting positive selection then descending ω values. Right panel, percentage of positively selected sites (by at least one method) over the length of the query coding sequence.

**Figure 4.**
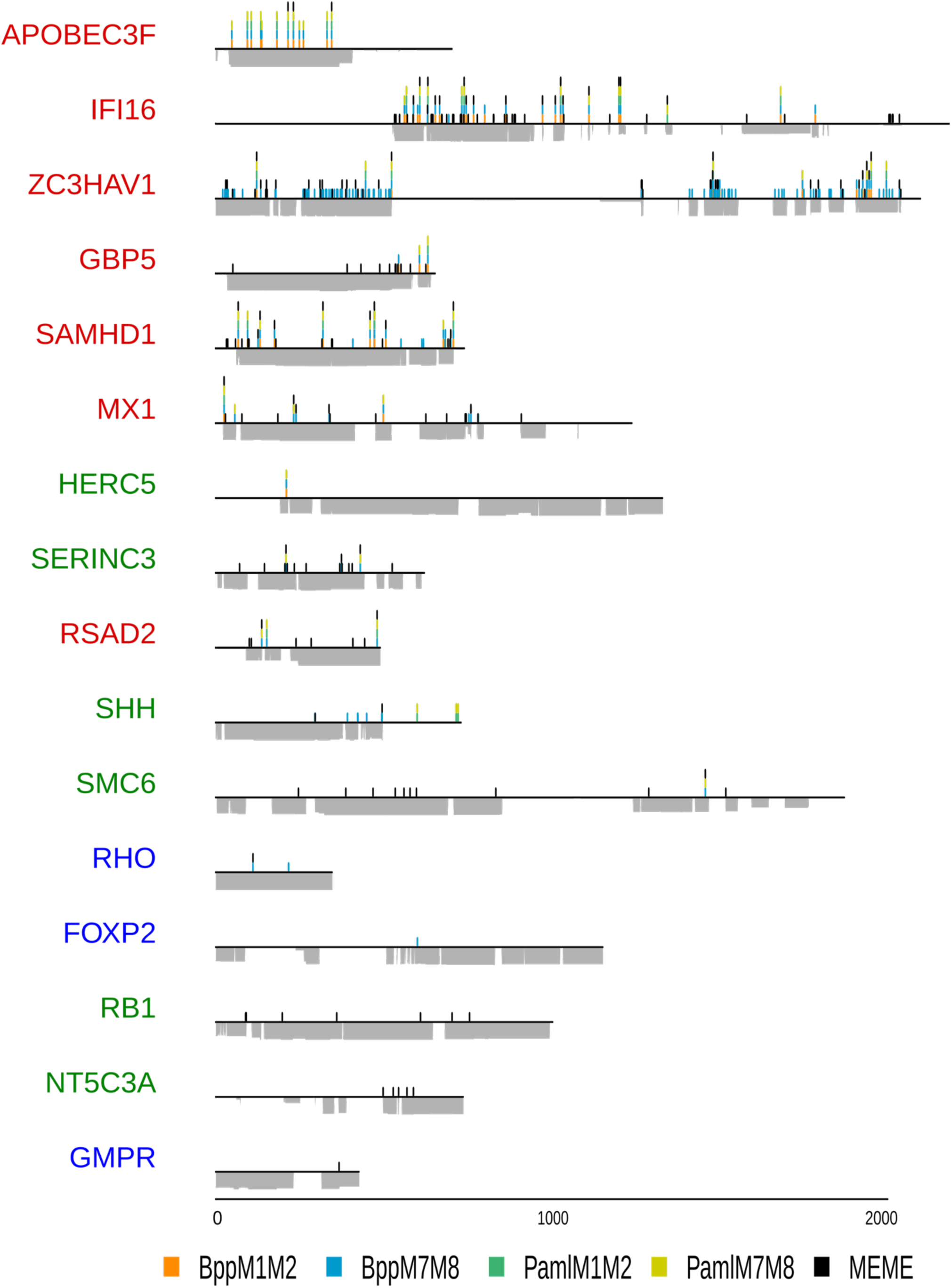
Positive selection patterns on nineteen primate genes. The nineteen genes studied are color-coded according to their selection profile category (Table 2) and follow the same order as in Figure 3. Genes without positively selected sites were excluded from this representation. Positively selected sites are represented as a spike (y axis) at their position along the alignment (x axis). Height of the peak is proportional to the number of methods that have identified the site as being under positive selection (posterior probabilities > 0.95 for Bio++ and PAML codeml, and p-value < 0.10 for MEME), with each method being represented by a different color (embedded legend). HYPHY MEME sites were only mapped if the gene was detected as under positive selection by BUSTED (p < 0.05). For each gene, the alignment coverage (number of sequences at a given position) is represented below the x axis.

### Validation dataset and method

To test our pipeline, we used a dataset of nineteen primate genes, for which evolutionary histories and positive selection profiles are either known and described in the literature or have been established within our laboratory in the past years (Table 2). We grouped those genes in three categories based on the clusters described in Murrell et al., 2016: “canonical arms-race genes” such as APOBEC3G and SAMHD1 (Table 2, red column), “genes described as presenting various selection profiles” (Table 2, green column), such as HERC5 or SERINC3, either regarding the methods employed to detect positive selection or the strength of the detected signal, and “genes under no positive selection pressure” such as GADD45A and RHO/rhodopsin (Table 2, blue column). The goal was to validate our automatic DGINN method using data and findings from highly hand-curated phylogenetic and evolutionary analyses, and if possible enrich them. To assess the pertinence of our detection of duplication events, we included nine genes belonging to multigene families (annotated with an asterisk in Table 2). A gene was considered as part of a multigene family if it had at least one paralogue with over 50% reciprocal identity amongst primates (according to Ensembl). A member of the *APOBEC3* gene family was also included as an extreme example of genes involved in virus-host evolutionary arms-races and that have undergone numerous genetic innovations (Nakano et al 2017, Etienne et al., 2015, Desimmie et al., 2014; Sawyer et al., 2004). Another example of multigene family member included is HERC5, which exhibits antiviral activity (reviewed in Kluge et al 2005) and described in the literature as evolving under positive selection (Woods et al., 2014). Given that in this latter case the analyses were performed on a limited number of primate species (seven species) and that this may conduct to a bias in the signature of positive selection, HERC5 was included in the “various” category rather than in the “canonical” one.

**Table 2.** Validation dataset of nineteen primate genes with various evolutionary histories. Genes are categorized according to their selection profiles as reported in the literature. Asterisks (*) denote a gene belonging to a gene family with at least one paralogue in primates presenting over 50% reciprocal identity with the query gene according to Ensembl.

| Canonical arms race genes |  | Variable signs of positive selection |  | No positive selection |  |
| --- | --- | --- | --- | --- | --- |
| **APOBEC3F \*** | Murrel *et al.*, 2016 | **HERC5** | Woods *et al.*, 2014 | **FOXP2 \*** | Murrel *et al.*, 2016 |
| **IFI16 \*** | Cagliani *et al.*, 2014 | **NT5C3A** | In house analysis | **GADD45A \*** | In house analysis |
| **GBP5 \*** | McLaren *et al.*, 2015 | **RB1** | Murrel *et al.*, 2016 | **GMPR \*** | In house analysis |
| **MX1 \*** | Mitchell *et al.*, 2012 | **SERINC3 \*** | Murrel *et al.*, 2016 | **ISG20** | In house analysis |
| **RSAD2/Viperin** | Lim *et al.*, 2012 | **SHH \*** | Murrel *et al.*, 2016 | **RHO** | Murrel *et al.*, 2016 |
| **SAMHD1** | Lim *et al.*, 2012,<br>Laguette *et al.*, 2012 | **SMC6** | Abdul *et al.*, 2018 | **TREX1** | In house analysis |
| **ZC3HAV1/ZAP** | Kerns *et al.*, 2008 |  |  |  |  |

The primate species tree used to assess for duplication events is based on the one established by Perelman et al. (2011) and updated by Pecon-Slattery (2014), with minor modifications: species’ names according to the six-letter naming system nomenclature that is used in DGINN (and is similar to UCSC genome’s nomenclature: the first three characters of the organism’s genus and species classification in the format gggSss; e.g. *Homo sapiens* becomes homSap), species names were updated (e.g. *Tarsius syrichta* was replaced by carSyr for *Carlito syrichta*), *Rhinopithecus bieti* (rhiBie) and *Rhinopithecus roxellana* (rhiRox) were added as the closest relatives of *Rhinopithecus brelichi* (rhiBre). This modified tree is available on DGINN’s Github (https://github.com/leapicard/DGINN/blob/master/examples/ex_spTree.tree).

### Reconstruction of the evolutionary history of primate Guanylate-binding protein (GBP) family

Homologs for human GBP4 and GBP6 were retrieved online through Blastn (https://blast.ncbi.nlm.nih.gov/) against the nr database limited to primates (taxid:9443). Sequences were manually selected to span as many primate species as available. Their accession numbers were added to the list of accession numbers previously obtained from the DGINN run from the human GBP5 query, then DGINN was run from the accession step to the duplication step (steps 2-5) to determine the new orthologous relationships and reconstruct the different gene trees.

### Resources

DGINN was run on the nineteen genes in a cluster environment (PSMN, http://www.ens-lyon.fr/PSMN/) in two stages. The first one ran from blast step against the NCBI non-redundant nucleotide *nr/nt* database circumscribed to primate species, with default settings (2 CPUs for each gene) until the identification of recombination events (steps 1-7, Figure 1). The second stage focused solely on positive selection analyses (step 8, 1 CPU for each alignment). Running times are summarized in Table 3.

**Table 3.** DGINN running times. For each gene, the running time of “Steps 1-7” (Figure 1) and “Step 8” is shown. Times for Step 8 (positive selection analyses) are only shown for the query genes of the validation dataset following attribution of orthologous groups (Table 4).

|  | **Steps 1 – 7** | **Step 8** |
| --- | --- | --- |
| **APOBEC3F** | 12:18:39 | 04:11:18 |
| **FOXP2** | 05:26:33 | 5 days, 23:28:04 |
| **GADD45A** | 01:24:26 | 02:17:59 |
| **GBP5** | 14:04:30 | 06:43:06 |
| **GMPR** | 03:51:52 | 07:50:42 |
| **HERC5** | 04:03:01 | 15:36:40 |
| **IFI16** | 05:45:16 | 5 days, 8:01:45 |
| **ISG20** | 01:34:50 | 08:01:08 |
| **MX1** | 02:34:42 | 1 day, 18:16:17 |
| **NT5C3A** | 00:53:48 | 4 days, 20:17:21 |
| **RB1** | 00:14:09 | 14:16:03 |
| **RHO** | 00:06:30 | 02:05:53 |
| **RSAD2** | 01:11:31 | 18:40:09 |
| **SAMHD1** | 00:51:44 | 3 days, 13:26:08 |
| **SERINC3** | 01:21:46 | 11:55:16 |
| **SHH** | 01:54:44 | 06:12:50 |
| **SMC6** | 02:48:41 | 2 days, 20:34:48 |
| **TREX1** | 01:01:04 | 09:43:10 |
| **ZC3HAV1** | 02:38:52 | 2 days, 13:27:12 |

### Availability

All scripts and documentation are freely available on Github (https://github.com/leapicard/DGINN) and as a Docker on DockerHub (https://hub.docker.com/r/leapicard/dginn). Example files to test DGINN are available to the users on GitHub. A specific script for the extraction of batch results, parseResults.py, is also available on the same Github. A graphical interface, which uses the file produced by parseResults.py as input and produces basic figures from the results (as in Figures 3-4), has also been developed and is available at https://rna-seq.shinyapps.io/DGINN_Pipeline_Visualization/. All results obtained and presented in this manuscript are available on the GitHub (https://github.com/leapicard/DGINN_validation).

## Results and Discussion

### 1 Presentation and novelties of the DGINN pipeline

The DGINN pipeline presents an end-to-end solution for the phylogenetic and automated detection of genetic innovations on protein-coding genes that are suspected to have undergone adaptive evolution. It automates the search for homologous sequences, their codon alignment and the reconstruction of phylogenetic histories. This is followed by the identification of marks of genetic innovations: (i) duplication events (also allowing for the identification of orthologous groups), (ii) recombination events (also limiting bias in subsequent positive selection analyses), (iii) positive selection through different methods.

The detailed presentation of the steps is found in the Method section.

Key novelties of the DGINN pipeline include a major focus on its flexibility of use: as such, it is possible to enter at any step in the pipeline without deep knowledge of the command line. The possibility to search within a single pipeline for diverse mechanisms of genetic innovations and using different methods for positive selection analyses translates to saved time compared to independent performance of each analysis. Moreover, though DGINN is designed to screen large datasets, it can also be used to perform gold-standard analyses on a single gene of interest with ease. For example, in the analyses of Lahaye et al (Lahaye et al., 2018), positive selection analyses on the *NONO* gene were performed through the use of DGINN to determine the evolutionary history of this newly discovered sensor of the Human Immunodeficiency Virus (HIV) capsid. Finally, DGINN includes key features detailed hereafter which are novel in such pipelines and allow for a more versatile use than just the detection of positive selection.

#### Automatic retrieval of homologous sequences and constitution of orthologous groups by tree reconciliation

The first important step for the identification of genetic innovations in a protein-coding gene is the retrieval of orthologous sequences of this gene, in as many species as possible in a given range, clade or family of interest to the user. Automating this step is a challenge as the evolutionary characteristics of orthologous genes vary a lot (between organisms, between copies in different species, according to different molecular clocks or environmental constraints). Usually, this step is time consuming and demands high manual curation. This is even more true for genes that have rapidly evolved. Most available tools for the detection of positive selection rely on user-provided alignments or are limited to fixed input species as PosiGene (Sahm et al., 2017). To circumvent these limits, DGINN uses BLAST against the NCBI online databases (see Methods – Steps 1-2). This approach makes the search for homologues simpler and relies on a widely-used and well-known tool, BLAST, which can be parameterized by the user. As true orthologous genes are identified through a subsequent reconciliation step, the user can cast a wide net by tuning parameters in terms of minimum coverage, e-value, identity, and species concerned.

From a set of homologous sequences, true orthologous groups are identified through a reconciliation software, Treerecs (Comte et al., 2019) and additional homemade scripts (Steps 3-5). Using tree reconciliation instead of annotations or tools such as OMA or Eggnogg (Altenhoff et al., 2018, Huerta-Cepas et al., 2016) may be particularly advantageous when working with non-model species, *unknown* genes, and recent duplication events. By separating the two phases of homology retrieval and orthology identification, we ensure that the user can change BLAST parameters without compromising the validity of the subsequent positive selection analyses.

#### DGINN detects gene duplication events, which may themselves be hallmarks of genetic innovation

While tools for the detection of positive selection abound, they often leave aside the detection of other hallmarks of genetic innovations, such as duplications (Daugherty and Zanders 2019). Very often, duplicated genes are even taken out of the analysis entirely to avoid bias during the detection of positive selection (Kosiol et al., 2008). However, this may lead to missing potential genes of interest and dismissing the gene copies that have been under adaptive evolution. On the contrary, DGINN looks for duplication events, as signals of potential genetic innovation as well as to identify relevant groups of orthology for further analyses. Similarly, tools which perform orthologous assignments from annotations cannot be trusted to detect either recent duplications or ancient ones on non-model species. To our knowledge this is the first time this feature is included in an automated pipeline searching for genetic innovation. The importance of accounting for those events is shown through the numerous genes involved in genetic conflicts which have undergone duplications and subsequent diversification (Daugherty and Zanders 2019). For example, many antiviral effectors, also called restriction factors, belong to multigene families, where duplicated copies have evolved varied antiviral functions and/or virus-host interfaces/determinants, such as the Mx (Myxovirus resistance) Dynamin Like GTPases Mx1 and Mx2 (Haller et al., 2015), the guanylate-binding proteins GBPs (Tretina et al., 2019, Huang et al., 2019), the primate *APOBEC3* gene family (Munk et al., 2012, Desimmie et al., 2014, Etienne et al., 2015, Nakano et al 2017) or the genes from the *TRIM* family (Malfavon-Borja et al., 2013).

#### Accounting for recombination allows for the detection of an important source of genetic innovation, while also avoiding bias in subsequent positive selection analyses

DGINN uses GARD to detect significant recombination breakpoints along the aligned sequences. As previously mentioned, recombination and gene conversion may be major sources of genetic innovations (in particular in the context of large gene families), and are widely ignored in existing pipelines. One example is the *TRIMcyp* gene present in certain primate species which results from the recombination and fusion of a *cypA* gene with the antiviral *TRIM5* gene leading to a change of antiviral specificity (Malfavon-Borja et al., 2013). Moreover, recombination may also itself bias phylogenetic reconstruction and positive selection analyses (Anisimova et al., 2003, Posada and Crandall, 2002), as exemplified by the multiple recombination and gene conversion events that occurred in the *Mx* gene family during mammalian evolution (Mitchell et al., 2015). To date, only PSP (Su et al., 2013) and PoSeiDon (Fuchs et al., 2017) pipelines account for such events in their workflow. In DGINN, detecting recombination events thus serves two purposes: identifying one possible hallmark of genetic innovation and avoiding bias in positive selection analyses.

#### DGINN integrates numerous methods for the detection of positive selection

The detection of signatures of positive selection is a key part of the pipeline. Indeed, very few pipelines include different models than the ones from PAML (Stern et al., 2007, Su et al., 2013). In DGINN, we decided to implement various methods with different underlying models, so the results obtained are more robust and can be balanced between methods. It also helps to “rank” the importance of signatures on genes when a large dataset is screened. The methods and models are described in the Method section, Step 8. In addition to the most used PAML codeml, we included Bio++ bppml with similar but non-stationary models. Of note, on our validation dataset, Bio++ bppml consistently performed better than PAML codeml when it comes to calculating likelihoods (Supplementary Table 1). Moreover, because of its versatility, Bio++ allows for more parameterization and the easy declaration of many modelings that would permit to detect positive selection under user-defined scenarios (e.*g.* using non-homogeneous mixture models, or other kinds of models such as allowing amino-acid specificity or simultaneous substitutions (Weber et al., 2019, Zaheri et al., 2014)).

Lastly, HYPHY is a good complement in those analyses, as shown in various studies (for example Malfavon-Borja et al., 2013, McBee et al., 2015, Rowley et al., 2016, Abdul et al., 2018, Schultz and Sackton, 2019). We thus decided to include two methods from the HYPHY package: one that considers the impact of positive selection at the level of the gene itself, using a branch-site model (BUSTED, Murrel et al., 2015), and another one which detects episodic positive selection at the site level (MEME, Murrel et al., 2012).

### 2 Validation

We tested our pipeline on nineteen primate genes selected for their various evolutionary histories and positive selection profiles (Table 2). These genes were grouped in three categories based on the clusters described in Murrell et al., 2016: “canonical arms-race genes” such as MX1 and SAMHD1, “genes described as presenting various selection profiles”, such as HERC5 or SERINC3 “genes under no positive selection pressure” such as GADD45A and RHO/rhodopsin (Table 2). The intermediate category was attributed on the basis of the methods employed to detect positive selection or the strength of the detected signal (see Method section).

#### An overview of the complete execution of DGINN on a protein-coding gene, HERC5

A brief overview of DGINN’s workflow on a specific gene, HERC5, is presented in Figure 2. The Blast search returned 71 primate homologous sequences, of which twelve were eliminated by the subsequent filters, yielding to a total of 59 sequences. As a duplication event was detected by Treerecs, these 59 sequences were then automatically (and correctly) split into two groups: one with 32 sequences corresponding to HERC5 and one with 27 sequences corresponding to HERC6. No recombination event was identified and the positive selection analyses then followed. All methods found highly significant evidence of positive selection on the complete alignment of 59 mixed HERC5-HERC6 sequences, with p-values ranging from 2.24e^-05^ to 2.27e^-13^ for PAML and Bio++ models. However, after separating the two paralogues into orthologous groups, it appeared that most of this signal was driven by the positive selection on HERC6 (p-values of 4.38e^-11^ to 8.10e^-15^ for PAML and Bio++ models), while the signal on HERC5 sequences was present but much more modest (p-values, 0.030 to 0.004), with BUSTED even returning a non-significant p-value for positive selection on that alignment. The positive selection results therefore highlight the necessity to properly separate paralogues from each other prior to performing the analyses. For a query on the *HERC5* gene, keeping the initial mixed alignment could have caused a mistaken conclusion that primate HERC5 has been under very strong positive selection, though the signal was mostly driven by HERC6. Moreover, the sites identified as under positive selection on that alignment would also be erroneous. Overall, the complete DGINN analyses with HERC5 as query took less than 20 hours (Table 3, 4h03 for the data mining and phylogenetics, and 15h36 for the detection of genetic innovations *per se*).

**Table 4.** Groups of orthologues reconstructed by DGINN, using long-branch partition and TreeRecs for identification of duplication events. For each gene of the validation dataset, are represented the orthologous groups that were identified, the number of sequences per group, the orthologues present in the group and the method used to separate the groups (long branch (LB) partition or TreeRecs-based). Groups kept for subsequent analyses are highlighted in yellow.

| Query | Group | Number of sequences | Gene | Type |
| --- | --- | --- | --- | --- |
| APOBEC3F | 1 | 11 | APOBEC3B | Treerecs-based |
|  | 2 | 56 | APOBEC3D + APOBEC3B |  |
|  | 3 | 16 | APOBEC3F |  |
|  | 4 | 94 | APOBEC3G |  |
|  | 5 | 10 | APOBEC3F |  |
| FOXP2 | 1 | 77 | FOXP2 | LB |
|  | 2 | 59 | FOXP1 |  |
| GADD45A | 1 | 30 | GADD45A | LB |
|  | 2 | 4 | GADD45B |  |
| GBP5 | 1 | 48 | GBP3 | Treerecs-based |
|  | 2 | 28 | GBP1 |  |
|  | 3 | 32 | GBP7 + GBP1 |  |
|  | 4 | 36 | GBP2 |  |
|  | 5 | 24 | GBP5 |  |
| GMPR | 1 | 104 | GMPR2 | LB |
|  | 2 | 34 | GMPR |  |
| HERC5 | 1 | 27 | HERC6 | Treerecs-based |
|  | 2 | 32 | HERC5 |  |
| IFI16 | 1 | 60 | IFI16 | Treerecs-based |
|  | 2 | 26 | MNDA |  |
| MX1 | 1 | 60 | MX2 | LB |
|  | 2 | 55 | MX1 |  |
| SHH | 1 | 25 | IHH | LB |
|  | 2 | 22 | SHH |  |
| TREX1 | 1 | 35 | TREX1 | LB |
|  | 2 | 14 | ATRIP |  |

#### Detection of ancestral duplications allows for proper assignation of orthologous groups

We identified genes as belonging to multigene families if at least one member had over 50% reciprocal identity with our gene query according to ENSEMBL annotations (Table 2). Given this definition, we were able to retrieve multiple family members for the majority of the genes belonging to such families, when performing BLAST with the minimum coverage (50%) and identity (70%) values. The sole exception was SERINC3, for which no paralogue was returned through our Blast search. Two additional exceptions were observed, first with HERC5, for which the Blast search also returned HERC6 sequences, though reciprocal identity between the two paralogues was below our threshold. The second case concerned TREX1, for which the Blast search also returned sequences annotated as ATRIP, an adjacent gene. Given that read-through transcription of TREX1-ATRIP occurs naturally and yields a non-coding transcript, it is probable that those sequences annotated ATRIP actually represents the non-coding transcript and not the mRNA of the ATRIP gene. This explains the retrieval of ATRIP-annotated genes through Blast despite the two genes not being strictly homologous.

DGINN efficiently reconstructed orthologous groups (Table 4). Indeed, in the case of multigene families (from two to five paralogues retrieved here), we were able to properly reconstruct orthologous groups for our genes of interest, without mixture with other paralogues. Our approach allowed us to separate the different family members retrieved through BLAST in groups which did not mix paralogous sequences through long branch partition (LB) and/or through reconciliation (Treerecs). For example, using the human CCDS sequence of FOXP2 as input in DGINN, we retrieved sequences from both FOXP2 and its paralogue FOXP1. The tree reconstructed from their alignment featured a branch over 50 times longer than the mean length of the tree’s branches, and by automatically splitting the sequences separated by that branch, we were able to reconstitute two groups corresponding to the paralogues. However, paralogues from other families may not have diverged enough for long branch partition to be able to properly discriminate them into different groups. We resolved those through TreeRecs, reconciling the tree obtained from the Blast-retrieved sequences with the primate species tree. This is the case, for example, of the immune sensor IFI16, which was properly assigned to a different group than MNDA through our Treerecs-based approach.

Non-annotated sequences (such as those referred as LOCXXX in databases) were also assigned to groups through this process, showing that this method of attributing orthologous relationships might help with non-annotated sequences in the databases.

Of our nineteen genes of interest, only one presented some inaccuracies in the distribution of sequences to orthologue groups. With an APOBEC3F query, DGINN erroneously divided APOBEC3F itself in two different groups (group 3 and 5, Table 4). By further analyzing all the retrieved paralogues, we observed two mixes: in the APOBEC3F query, group 2 contained APOBEC3D and APOBEC3B sequences and APOBEC3B was split in two groups, and a similar pattern occurred in the GBP5 query, with GBP1 in groups 2 and 3 (Table 4). These errors could be explained by the particularly complicated evolutionary histories of those two expanded gene families during primate evolution (Münk et al., 2012, Desimmie et al., 2014, Nakano et al., 2017). This highlights a need to improve the management of the detection of duplication events in further versions of DGINN. Importantly, because such genes would be tagged by DGINN with “detected duplication events”, these cases would anyway not be missed by the user and the gene of interest could be reanalyzed through DGINN after curation.

#### Using several positive selection methods together allows for more sensitivity and specificity and a “ranking” of genes’ positive selection status during screening

Positive selection results were analyzed according to two different aspects. The first aspect focused on how many methods found a gene with significant evidence of positive selection (Figure 3, left panel – produced using the Shiny app openly available). The methods considered at this point were those on which a LRT could be performed: HYPHY BUSTED, the M1 *vs* M2 and M7 *vs* M8 models of PAML Codeml, and the M1^NS^ *vs* M2^NS^ and M7^NS^ *vs* M8^NS^ models of Bio++ bppml. Genes were ranked according to the number of positive results. This allowed us to compare the results obtained for the three categories of genes (Table 2). The canonical arms-race genes were all detected under positive selection by all five methods, with the exception of *RSAD2* which was detected by four methods (Figure 3). Genes which presented variable signs of positive selection in the literature (green category, Table 2) also fell into a middle category in the DGINN screen. Genes without signs of positive selection in previous studies (blue category, Table 2) displayed low signs of positive selection: detected by less than two methods in DGINN. Two genes were detected by two methods: *FOXP2* and *RHO. FOXP2* was detected by both PAML M7 *vs* M8 and Bio++ M7^NS^ *vs* M8^NS^, but both the mean omega and the very low number of sites detected under positive selection (n=1) suggested artefactual results. Similarly, *RHO* was detected by BUSTED and Bio++ M7^NS^ *vs* M8^NS^, but only two sites were detected. Therefore, our DGINN screen efficiently recapitulated results from published studies.

These results further highlight the advantage of using different methods within a single analysis to confirm results and discriminate for false positives. Doing this validation also showed that amongst those methods, BUSTED and PAML Codeml M7 *vs* M8 appeared the least conservative methods to detect positive selection.

The second aspect taken into account focused on the percentage of positively-selected sites. Overall, the arms-race genes displayed higher proportions of positively selected sites (2.4%-14.4%) compared to other genes (Figure 3, right side). However, this does not represent a hard rule, as some of those arms-race genes show rather low percentages, such as MX1 (around 2.4%). This suggests that ranking genes by the number of significant methods rather than the proportion of positive selection sites, as in Figure 3, is a better proxy for positive selection status.

#### DGINN recapitulates and expands the findings from previously published profiles of positively selected sites along genes

To identify the domains that have evolved under positive selection, we mapped every positively selected site detected by DGINN by a peak along the alignment (Figure 4, using the Shiny app). We further represented the height of the peak proportional to the number of methods detecting that site under significant positive selection, amongst five methods, M2 and M8 results of PAML codeml, M2^NS^ and M8^NS^ results of Bio++ bppml and HYPHY MEME (Figure 4). Overall, we observed similar patterns as described in the literature, especially on the canonical arms-race genes. For example, in the case of SAMHD1, we found most positively selected sites at the N- and the C-termini (Figure 4). This is in accordance with the findings that the N-ter and C-ter domains both play a role in the antiviral/escape determinants of primate SAMHD1 and that rapid evolutions at these sites are certainly adaptive as a result of lentiviral selective pressure (Fregoso et al., 2013, Lim et al., 2012, Laguette et al., 2012). In the case of *ZC3HAV1*/ZAP, we found the positively selected sites cluster at both extremities of the alignment (Figure 4). However, the middle portion without positively selected sites corresponds to a gap-enriched region in the alignment linked to the different possible isoforms of the gene. Interestingly, this shows that the maintenance of these gap regions in the alignment did not lead to an excess of false positive detection in DGINN. If we now consider the main ORF (with the gap-enriched region ignored), it appears that the positively selected sites are spread over the whole length of the gene. Previous results from Kerns et al., 2008 established that the C-ter domain in particular was under significant positive selection (Kerns et al., 2008). In contrast, the N-ter domain was not detected, probably because we used more methods and had more species/sequences available for analyses.

The differences between our results and the published ones for *APOBEC3F* (Murrell et al., 2016) were mainly due to the sequences used for the positive selection analyses. Indeed, our analyses excluded four hominoid species that were correctly retrieved in the early steps of DGINN but were erroneously assigned by TreeRecs to another group. The detection of positive selection was therefore only performed on a subset of primate sequences, spanning solely Old World monkeys. However, we have included the solution to such problems in DGINN thanks to its high flexibility. The user may retrieve the gene sequences (here *APOBEC3F*) from the different groups and re-enter DGINN at step 3/alignment (Figure 1 and Table 2) to obtain the complete evolutionary history and positive selection analyses.

For *MX1*, we were first surprised that we did not detect such a high signal of positive selection in the L4 loop as described in Mitchell et al., 2012. However, we found that this was mainly due to differences in the alignments, because PRANK (as opposed to ClustalX used in Mitchell et al., 2012) introduced many gaps in the L4 loop region due to the extremely-high divergence of the region. Whether MX1 adaptation to viral countermeasures has occurred by accumulation of non-synonymous changes and/or by indels in the L4 loop remains to be determined.

In the case of HERC5, four methods detected the gene as under positive selection during primate evolution (Figures 2-3), but only one site was identified as positively selected (Figure 4). These results differ from the ones reported in Woods *et al.*, 2014, who found a much larger number of residues under positive selection (n=50). This discrepancy, however, can be explained by the fact that Woods *et al.* identified positive selection on an alignment that included six non-primate species and only seven primate species, while ours focused exclusively on primates and included twenty species. It is therefore possible that a stronger selective pressure has occurred in placental mammals outside of primate evolution. Interestingly, in DGINN, our Blast search with HERC5 as query also automatically retrieved HERC6 sequences (Figure 2). The latter were then correctly assigned to a different orthologous group than HERC5. As previously reported (Paparisto et al., 2018), we identified strong evidence of positive selection on HERC6 (with five methods, Figure 2). This could mean that while both HERC5 and HERC6 have been evolving under positive selection in mammals, they have been subjected to different evolutionary constraints in primates, with a lower selective pressure on primate HERC5 *vs* HERC6. It further shows that DGINN is an efficient tool to screen not only the query genes but also the evolutionary history of their closest gene relatives that may have themselves be subjected to positive selection and would otherwise be missed by most analyses.

#### Identification of the loss of GBP5 during primate evolution using DGINN

The positive selection results obtained through DGINN screening for *GBP5* showed strong positive selection (identified by five methods). This was in accordance with previous results from McLaren et al., 2015. By analyzing the phylogenetic tree generated by DGINN for all the homologs retrieved with the *GBP5 query* (after step 4, Figure 5A), we found that no sequence from Old World monkeys were retrieved for *GBP5* through our Blast search. This absence was confirmed in the tree reconstructed with only *GBP5* sequences after orthologue group attribution (step 5, Figure 5B). However, (and as expected), the entire *GBP* gene family was not retrieved by DGINN using human GBP5 as query (with blastn 70% identity and 50% coverage); in particular, *GBP4* and *GBP6* were too divergent to be retrieved by DGINN. To reconstruct to *GBP* family evolutionary history, we independently retrieved primate sequences of *GBP4* and *GBP6* by blastn and added the new sequences to a large *GBP* family sequence file. This served as input to DGINN steps 2-5 to automatically perform alignments, phylogenies, and duplication/orthologous group detection. The final tree confirmed that *GBP5 is* absent in Old World Monkeys (Figure 5C). This might also be the case for GBP4, for which we did not retrieve sequences from Old World Monkeys; with the exception of two sequences from *Papio anubis* and *Mandrillus leucophoeus* that were annotated as “GBP4” but did not follow a typical orthologous phylogeny and branched more closely with GBP7 in our phylogeny (Figure 5C). Genomic analyses of the *GBP* locus in several primates confirmed that *GBP5* has been lost in the ancestor of Old World Monkeys during primate evolution, and that it may also be the case for GBP4 (Figure 5D). To explain our retrieval of the two sole Old Word Monkeys sequences, and their position in the phylogeny, one hypothesis could be that GBP4 has indeed been lost at a similar point in primate evolution than GBP5, and was then regained in some Old World monkey species through a duplication of GBP7. Overall, these results show that *GBP5* has been subjected to strong positive selection during primate evolution but has also entirely been lost in the *Cercopithecinae*. Whether part of this has been driven by pathogens such as lentiviruses (Krapp et al. 2016) or bacteria (Kim et al. 2012) should be investigated.

**Figure 5.**
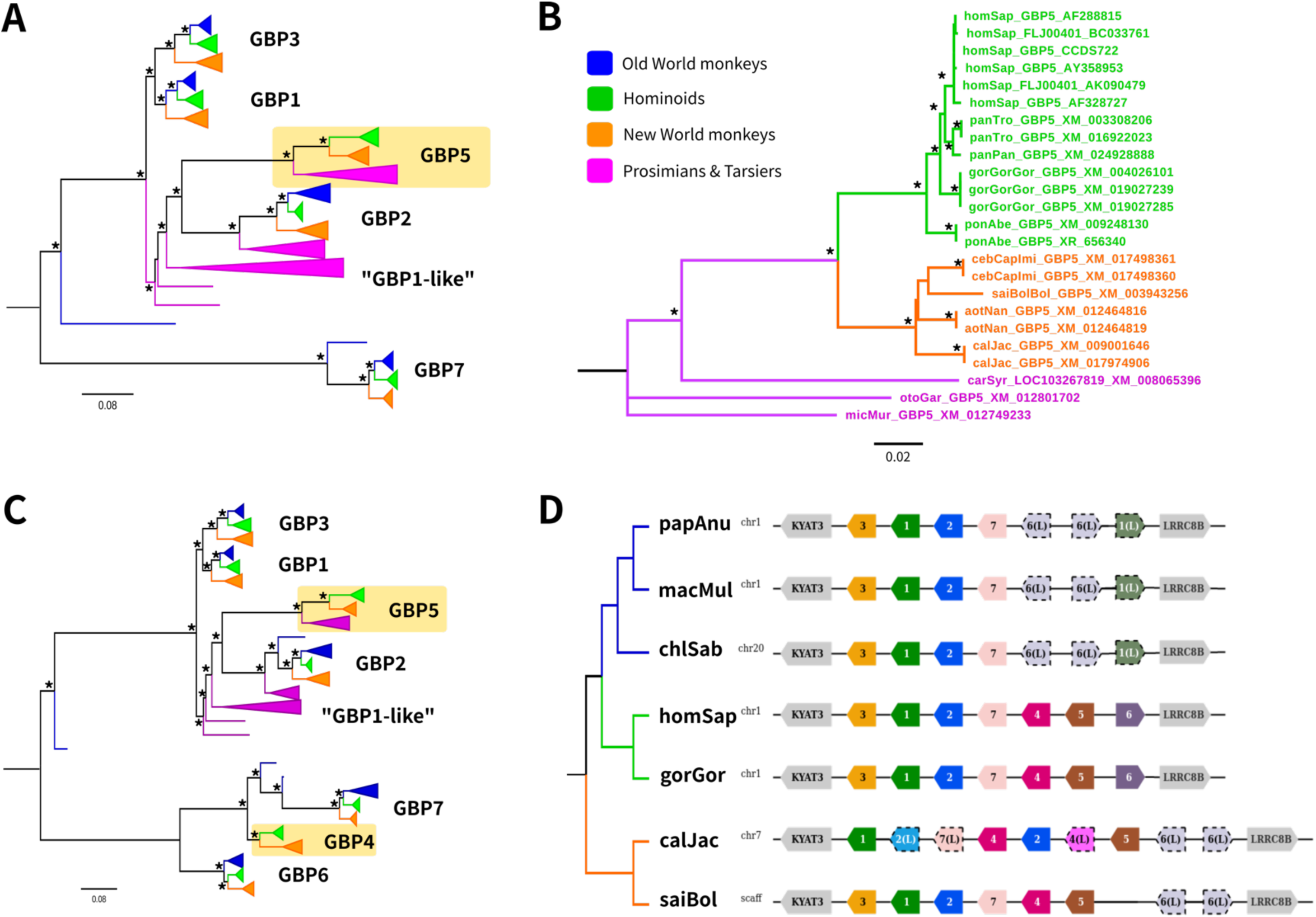
Evolutionary history of the primate GBP family. (A) Maximum-likelihood phylogeny established through DGINN based on a run on the GBP5 query (step 4). The four main primate lineages are identified by color-coding (embedded legend). Asterisks (*) denote nodes that are statistically supported by aLRT > 0.90. The *GBP5* group, which lacks Old World monkey sequences, is boxed in yellow. The scale bar represents the number of nucleotide substitutions per site and the tree was midpoint rooted. (B) Maximum-likelihood phylogeny of the *GBP5* group of primate orthologues established through DGINN screen (step 7). (C) Maximum-likelihood phylogeny of the whole *GBP* family performed in DGINN after manual addition of primate *GBP4* and *GBP6* sequences. (D) Diagram of the genomic locus of the *GBP* gene family in seven simian primate species. The reference genomes from the NCBI used were: papAnu (*Papio anubis*): Panu_3.0, macMul (*Macaca mulatta*): Mmul10, chlSab (*Chlorocebus sabaeus*): Chlorocebus_sabeus 1.1, homSap (*Homo sapiens*): GRCh38.p13, gorGor (*Gorilla gorilla*): gorGor4, calJac (*Callithrix jacchus*): Callithrix jacchus-3.2, saiBol (*Saimiri boliviensis*): saiBol1.0. Gene annotations and predictions are from the NCBI database. “X(L)” annotations with dotted outlines, such as 6(L), represent genes for which the orthology and paralogy relationships have not been determined. Alignments and phylogenies for panel A, B, and C can be found at https://github.com/leapicard/DGINN_validation/tree/master/GBPfamily/ (referred as 5A_aln, 5A_tree, etc.).

## Conclusion

We have developed DGINN, an integrative pipeline for the automatic detection of genetic innovations, and made it freely available through both GitHub and Docker. DGINN was validated for screening usage against nineteen primate genes. It automates and streamlines those analyses, allowing the user to simply provide the coding sequence of their gene of interest and a parameter file to complete the whole workflow, from retrieval of homologous sequences to the detection of orthology relationships, recombination events and positive selection.

Through our validation, we confirmed and expanded on results previously established in the literature. Genes described as engaged in arms-races with viruses were found under strong positive selection by all five methods included in DGINN. Our analyses allowed us to establish clearer profiles for the genes belonging to the “varied” category, owing to our inclusion of different methods for positive selection: this way, we were able to establish that some genes previously thought to present moderate signs of positive selection presented stronger signs than suspected. Little evidence of positive selection was found on the genes belonging to “no positive selection” category, in accordance to the literature.

An important feature of DGINN is its flexibility, which allows usage beyond its screening capacity. Indeed, in cases of dubious results, the possibility remains for the user to curate their input files and perform the appropriate analyses by entering DGINN at any of the downstream steps. This also means that the “positive selection” part might be of primary interest to scientists wishing to perform gold-standard positive selection analyses on their favorite gene, because they could enter their curated alignment and phylogeny and obtain results of positive selection analyses from five methods in a single query.

Using DGINN to analyze nineteen primate genes also allowed us to enrich some findings, notably on the importance of detecting duplications and properly ascribing orthologue groups, as exemplified by the case of HERC5 and its paralogue HERC6 in primates. The ability to check multiple members of a query’s gene family is a major advantage of DGINN, as it may allow the user to identify genes bearing signs of genetic innovations that they would not have analyzed otherwise. Improving the constitution of orthologue groups will remain an objective in future versions of DGINN.

## Acknowledgements

We thank Stéphanie Jacquet for her comments on the manuscript. We also thank Bastien Bousseau, Marie Cariou, Hélène Dutartre, Laurent Modolo, Xavier Morelli, and Guy Perrière for helpful discussions on this project. We gratefully acknowledge support from the PSMN (*Pôle Scientifique de Modélisation Numérique*) of the ENS de Lyon for the computing resources, and the PRABI (*Pôle Rhône-Alpes de BioInformatique*) for further bioinformatics support.

We thank all the contributors of publically available genome sequences, as well as the scientists who developed the methods included in DGINN.

This work was funded by the ANR LABEX ECOFECT (ANR-11-LABX-0048 of Université de Lyon, within the program “Investissements d’Avenir” (ANR-11-IDEX-0007) operated by the French National Research Agency) to LE and LG. LE is supported by the CNRS and by grants from the amfAR (Mathilde Krim Phase II Fellowship #109140-58-RKHF), the “Fondation pour la Recherche Médicale” (FRM “Projet Innovant” #ING20160435028), the FINOVI (“recently settled scientist” grant), the ANRS (#ECTZ19143, #ECTZ118944), and a JORISS incubating grant. LG is supported by the Université Claude Bernard Lyon 1 and the Swedish Center of Advanced Study. AC is supported by the CNRS and by grants from the ANRS, Sidaction and the ENS-L.

## Author Contributions

Conceptualization and Supervision: LE, LG

Formal analysis: LP, LE, LG

Pipeline development: LP

Funding acquisition: LE, LG

Investigation: LP, QG, OA, AC, LG, LE

Methodology: LP, LG, LE

Project administration: LE, LG

Resources: AC, LE, LG

Writing – original draft: LP, LE, LG

Writing – review and editing: All the authors

## Supplementary Information

**Supplementary Table 1.** Log likelihoods calculated by BIO++ and PAML codeml for each of the different models.

| File | Method | M1 | M2 | M7 | M8 |
| --- | --- | --- | --- | --- | --- |
| APOBEC3F | BPP | -2507,37 | -2491,43 | -2511,01 | -2491,47 |
|  | PAML | -3945,63 | -3930,68 | -3945,88 | -3930,68 |
| FOXP2 | BPP | -3997,30 | -3995,85 | -3999,96 | -3995,98 |
|  | PAML | -5860,15 | -5858,43 | -5861,65 | -5858,44 |
| GADD45A | BPP | -1239,99 | -1239,99 | -1241,28 | -1238,69 |
|  | PAML | -1291,75 | -1291,75 | -1292,06 | -1289,64 |
| GBP5 | BPP | -6138,60 | -6131,37 | -6139,83 | -6131,78 |
|  | PAML | -6516,42 | -6504,84 | -6518,34 | -6505,90 |
| GMPR | BPP | -3027,30 | -3027,30 | -3027,12 | -3025,84 |
|  | PAML | -3425,82 | -3425,82 | -3423,78 | -3423,53 |
| HERC5 | BPP | -10433,56 | -10430,06 | -10434,33 | -10430,37 |
|  | PAML | -11950,25 | -11945,50 | -11951,66 | -11946,35 |
| IFI16 | BPP | -11099,74 | -11075,12 | -11103,87 | -11075,78 |
|  | PAML | -18264,89 | -18223,86 | -18270,40 | -18225,35 |
| ISG20 | BPP | -1837,14 | -1837,14 | -1836,69 | -1836,69 |
|  | PAML | -3048,57 | -3048,57 | -3047,88 | -3047,42 |
| MX1 | BPP | -8537,86 | -8532,47 | -8543,10 | -8531,57 |
|  | PAML | -11329,45 | -11316,93 | -11335,61 | -11317,74 |
| NT5C3A | BPP | -2324,85 | -2324,85 | -2324,19 | -2323,64 |
|  | PAML | -4483,43 | -4483,43 | -4482,88 | -4482,16 |
| RB1 | BPP | -6873,89 | -6873,89 | -6874,36 | -6872,91 |
|  | PAML | -7244,38 | -7244,38 | -7242,93 | -7242,93 |
| RHO | BPP | -2910,93 | -2910,93 | -2913,24 | -2908,65 |
|  | PAML | -2967,07 | -2967,07 | -2953,69 | -2953,12 |
| RSAD2 | BPP | -4248,12 | -4248,12 | -4248,75 | -4243,00 |
|  | PAML | -4875,51 | -4868,59 | -4874,28 | -4865,51 |
| SAMHD1 | BPP | -7522,22 | -7495,71 | -7534,66 | -7497,30 |
|  | PAML | -8219,63 | -8186,06 | -8225,38 | -8187,80 |
| SERINC3 | BPP | -4751,87 | -4749,23 | -4755,82 | -4749,69 |
|  | PAML | -5373,70 | -5370,57 | -5376,40 | -5371,08 |
| SHH | BPP | -3576,79 | -3576,79 | -3580,07 | -3575,89 |
|  | PAML | -4855,15 | -4841,25 | -4853,34 | -4837,51 |
| SMC6 | BPP | -8971,83 | -8971,83 | -8974,05 | -8970,65 |
|  | PAML | -12342,77 | -12342,77 | -12341,63 | -12336,91 |
| TREX1 | BPP | -3421,42 | -3421,42 | -3421,35 | -3421,15 |
|  | PAML | -5080,28 | -5080,27 | -5080,76 | -5080,15 |
| ZC3HAV1 | BIO++ | -12413,91 | -12384,63 | -12418,39 | -12392,87 |
|  | PAML | -17617,28 | -17585,61 | -17619,80 | -17585,77 |

